# treecompareR: Tree Visualizations of Chemical Space

**DOI:** 10.1101/2025.04.04.647092

**Authors:** Paul Kruse, Caroline Ring

## Abstract

This paper presents the **treecompareR** package for `R`, which provides tools for reproducible visualizations of data through the use of taxonomies. The package builds on developments from **ggplot2** and **ggtree** to provide visualizations tailored for use with taxonomic classification data. Additionally, it provides tools that leverage developments in network analysis to compare data sets. While designed specifically for chemical classification objectives using ClassyFire (http://classyfire.wishartlab.com/), **treecompareR** also provides tools for use with more general taxonomies. **treecompareR** is available from GitHub at https://github.com/USEPA/treecompareR.

## 1. Introduction

Classification of objects has been a source of much development of organizational structures across various fields of science. Organizing objects into a coherent structure helps one to understand how these objects relate to each other and can lead to the discovery of connections between various sets of objects. How one organizes the objects may depend on underlying attributes that the objects possess.

Ontologies provide a framework for comparing objects through a variety of different relation types. These relation types create a network structure that can be studied using graph theory techniques. One of the more actively developed ontologies in recent science is the Gene Ontology, which aims to develop a computation model of biological systems. The ontology considers the three domains defined as molecular function, cellular component, and biological process with relationships between terms within each of these domains and between the three domains as well. Using these domains, the ontology describes genes and how they relate to various areas of interest in contemporary biological research.

Within the universe of ontologies sits the set of organizing structures known as taxonomies. Restricting the attention to taxonomies, one can focus specifically on the “is a” relation, e.g. “a dog (canis familiaris) is a canis” and study how objects relate to one another under this specific relationship scheme. One well known example of this organizational structure in use is the Linnaean system for classifying life, sometimes referred to as the “tree of life”. As a taxonomy, the organizational structure is given by a tree, which consequently lends itself to analysis using tree-based techniques from graph theory.

In 2016, the ClassyFire (http://classyfire.wishartlab.com/) tool was introduced, in the work of Djoumbou Feunang, Eisner, Knox, Chepelev, Hastings, Owen, Fahy, Steinbeck, Subramanian, Bolton *et al*. (2016). This work presented a chemical ontology, ChemOnt, consisting of 4825 classification labels. These labels are organized in a taxonomy structure with pairs of distinct labels either satisfying a simple “is a” relation or not. The ClassyFire tool takes in chemical identifiers such as InChIKey or SMILES strings and provides a classification of the chemical within the taxonomy. In this manner, one can examine a data set consisting of chemical data from the viewpoint of where the chemicals lie within the ChemOnt taxonomy structure.

In the study of cheminformatics, machine-learning models such as Quantitative structure-activity relationship (QSARs) are constructed to link physical and structural properties with biological interactions. For these models to produce accurate and reliable predictions, an analysis of the training data and targeted use data is essential. This consideration of the domain of applicability of the model is important for determining when the trained model’s predictions can be considered reliable given an input chemical. In many cases, this analysis can be carried out using model descriptors, for example by using structural descriptors of the molecules used for training the model. However, while descriptors may accurately describe the domain of applicability, they often can be hard to interpret and they do not necessarily lend themselves that well to visualization.

Understanding and overcoming these issues in one’s ability to interpret the data are especially important when comparing pairs of data sets, such as training and test sets or training and prediction sets. While it may be feasible to examine how each individual data set is characterized by model descriptors, how they overlap requires additional consideration. Is the overlap broad or does it focus on specific areas? Are there gaps between the training data and prediction data that may reduce the confidence users have in model predictions? Answering these and other questions addresses concerns of model applicability to data sets in general and how the model can be improved regarding gaps in training data compared to prediction or test data.

One approach to addressing these naturally arising concerns is through visualizations that can increase the user’s ability to interpret the data sets. Since the ClassyFire ChemOnt ontology is organized in a taxonomy structure, one can leverage the tree structure not only for quantitative analysis but also through the use of tree visualizations. These visualizations can reveal the extent to which there are overlaps within the data and the degree of such overlaps. The diagrams also show very clearly when there are gaps between the data sets and broad coverage, or lack thereof, within the chemical universe as it is represented by ChemOnt.

However, such visualizations are limited in terms of what is currently available. One may use the **ggplot2** `R` package to visualize statistics based on the taxonomic information corresponding to classifications of a data set Wickham (2016). But **ggplot2** is a very broad package and not designed for handling chemical classification data in particular. Since the ChemOnt taxonomy is given by a tree structure, one can use tree visualization packages such as **ggtree** to visualize this structure Yu, Smith, Zhu, Guan, and Lam (2017). However, the **ggtree** package was designed with phylogenetic trees in mind, which are generally presented as rooted binary trees (possibly with additional data). The ChemOnt ontology is neither a binary tree nor does it carry additional explicit data, so **ggtree** is again a very broad approach to this. The package **treecompareR** leverages several `R` packages designed for tree manipulation and exploration, including **phytools** Revell (2024), **phangorn** Schliep (2011); Schliep, Potts, Morrison, and Grimm (2017), **Ape** Paradis and Schliep (2019), **Treeio** Wang, Lam, Xu, Dai, Zhou, Feng, Guo, Dunn, Jones, Bradley, Zhu, Guan, Jiang, and Yu (2020). It also combines tree visualizations with heatmaps, in particular using the package **ComplexHeatmap** Gu, Eils, and Schlesner (2016).

We present the package **treecompareR** as a solution to the specific needs of creating a pipeline to classify chemicals in a data set, visualize the chemical space they represent, and compare the data sets from the point of view of chemical space. We introduce the package through a case study analyzing two specific data sets. We characterize two data sets, named `BIOSOLIDS` and `USGSWATER`, that were collected from the CompTox Chemical Dashboard Williams, Grulke, Edwards, McEachran, Mansouri, Baker, Patlewicz, Shah, Wambaugh, Judson *et al*. (2017). The `BIOSOLIDS` data describe chemicals detected in biosolids, as reported by three national sewage sludge surveys and eight biennial reviews, while the `USGSWATER` data describe chemicals found in water by the USGS.

We are interested in studying these two data sets together for a variety of reasons. The presence of chemicals detected in water is important information to consider when determining risk assessment for human and ecological health. Not only does this indicate what chemicals human activity contributes to water, but also what to expect to be present in water that is used in commercial, industrial, and residential settings. Chemicals detected in biosolids indicate both the chemicals that are already present in water before being used, and the chemicals that human activity contributes to water and persists through the treatment process of waste water. By examining these two sets, we can get a better idea of which chemicals are introduced directly through human activity and use, and which chemicals are already in the background water as well as those that may be successfully removed from the water and captured within the biosolids.

## 2. ClassyFire

In this paper, we use the `BIOSOLIDS2021` and `USGSWATER` data sets, which are available for download from the US Environmental Protection Agency’s Comptox Chemicals Dashboard https://comptox.epa.gov/dashboard/chemical-lists. Note, the names of these data sets as listed on the CompTox Dashboard Chemical List are `BIOSOLIDS2021` and `USGSWATER`, respectively.

These data sets include a list of chemicals, relevant identifiers such as `DTXSID`, `InChIKey`, `SMILES`, physico-chemical property data such as `MOLECULAR_FORMULA`, `AVERAGE_MASS`, and `MONOISOTOPIC_MASS`, and publication and quality data such as `PUBCHEM_DATA_SOURCES` and `QC_Level`.

Before we start comparing these data sets, we must provide each chemical in each data set classification data from ClassyFire. To do this, we use the classification pipeline in **treecompareR**, construct a tree that encodes the taxonomy used in this classification scheme, and use this tree for data analysis and to create visualizations.

To achieve these aims, we use the ClassyFire tool found at http://classyfire.wishartlab.com/. ClassyFire uses the ChemOnt ontology, consisting of 4825 labels over 11 levels of classification, to classify chemicals. The ontology is given a taxonomic structure, that of a tree, with each label given a unique parent label and potentially having several children labels. This relationship data between labels can be found at the ChemOnt Taxnodes page http://classyfire.wishartlab.com/tax_nodes.json in JSON format. To construct a tree that encodes this structure, we must first download this data or use the file `chemont_parent_child.rda` from **treecompareR** with column names `Name`, `ID`, `Parent_ID`. With this data in hand, we construct the tree as a `phylo` object as follows.

~~~
*R> chemont_taxonomy <- generate_taxonomy_tree(tax_nodes = chemont_df)*
*R> sapply(chemont_taxonomy*,
*+        function(x) head(x)*,
*+        simplify = FALSE*,
*+        USE*.*NAMES = TRUE)*
$edge
     [,1] [,2]
[1,] 3613 3614
[2,] 3614 3615
[3,] 3615    1
[4,] 3615    2
[5,] 3615 3616
[6,] 3616 3617
$Nnode
[1] 1213
$tip.label
[1] “Thiepanes”                     “Benzoxazolines”
[3] “Polychlorinated dibenzofurans” “Polybrominated dibenzofurans”
[5] “Phthalic anhydrides”           “Pterofurans”
$edge.length
[1] 9.090909 11.363636 79.545455 79.545455 19.886364 19.886364
$node.label
[1] “Chemical entities”            “Organic compounds”
[3] “Organoheterocyclic compounds” “Benzofurans”
[5] “Dibenzofurans”                “Chlorinated dibenzofurans”
~~~

The output of the `generate_taxonomy()` function is a list of two objects, the `phylo` object representing the tree, and a data frame used in creating the `phylo` object. Observe the structure of these in the code chunk above.

The `phylo` object itself is a list, describing the edges of the tree, the number of internal nodes, the tip labels and internal node labels, and a set of edge lengths generated in the construction of the tree. One thing to note is that the edge lengths are artificial and are only used to speed up the creation of the tree, but do not represent any actual data corresponding to the tree. However, we retain the edge lengths as they are useful in creating visualizations.

To construct a tree for other taxonomies, one can enter in a path to a JSON file or an enter in a data frame. The JSON file must have three columns, the first the name of the label `Name`, the second its identifier `ID`, and the third the parent identifier `Parent_ID`. For the root, the parent ID should be `NA`. If the data is in the form of a data frame, it must be organized in the same way the JSON file is, with column names given by the same column names in the same order.

Next, we use the ClassyFire tool to provide classifications for the chemicals in our data sets. To simplify this process, **treecompareR** has a data pipeline that allows for programmatic acquisition of this data using the ClassyFire API. For chemicals missing certain identifying data such as InChIKey or SMILES string data, we can use functions from the `R` package **ctxR** Kruse, Ring, Feshuk, Brown, and Thunes (2024). We demonstrate how to use these below.

~~~
*R> CASRN <- c(‘117964-21-3’, ‘11006-76-1’)*
*R> chemical_search <- ctxR::chemical_equal_batch(word_list = CASRN) R> chemical_ids <- ctxR::get_chemical_details_batch(*
*+   DTXSID = chemical_search$valid$dtxsid*
*+)*
*R> data*.*table::setnames(x = chemical_ids*,
*+                       old = ‘inchikey’*,
*+                       new = ‘INCHIKEY’)*
*R> chemicals_class <- classify_datatable(*
*+   chemical_ids[*, .*(INCHIKEY, dtxsid, casrn)]*
*+)*
*R> str(chemicals_class)*
Classes ‘data.table’ and ‘data.frame’: 2 obs. of 17 variables:
$ INCHIKEY   : chr NA “AAFUUKPTSPVXJH-UHFFFAOYSA-N”
$ dtxsid     : chr “DTXSID40880080” “DTXSID9074775”
$ casrn      : chr “11006-76-1” “117964-21-3”
$ smiles     : chr NA “BrC1=CC(Br)=C(Br)C(Br)=C1OC1=C(Br)C(Br)=C(Br)C=C1Br”
$ inchikey   : chr NA “InChIKey=AAFUUKPTSPVXJH-UHFFFAOYSA-N”
$ report     : chr NA “ClassyFire returned a classification”
$ kingdom    : chr NA “Organic compounds”
$ superclass : chr NA “Benzenoids”
$ class      : chr NA “Benzene and substituted derivatives”
$ subclass   : chr NA “Diphenylethers”
$ level5     : chr NA “Bromodiphenyl ethers”
$ level6     : chr NA NA
$ level7     : chr NA NA
$ level8     : chr NA NA
$ level9     : chr NA NA
$ level10    : chr NA NA
$ level11    : chr NA NA
attr(*, “.internal.selfref”)=<externalptr>
attr(*, “sorted”)= chr “INCHIKEY”
~~~

In this example, we input into the `ctxR::chemical_equal_batch()` function the list of CASRN strings. This searches and returns chemicals that match these values exactly. Next, the function `ctxR::get_chemical_details_batch()` uses the `dtxsid` column values to retrieve the chemical details for these. Finally, we input as a `data.table` with columns `INCHIKEY`, `dtxsid`, and `casrn` into the function `classify_datatable()`, which hits the ClassyFire API and attaches the retrieved information to the input data, as displayed. Note that the `CASRN` with value ‘117964-21-3’ has both an `inchikey` and `smiles` value while the `CASRN` with value ‘11006-76-1’ has neither piece of information. The ClassyFire data is available for the first chemical, but not the second.

**Table 1:** Overview of data contained in the usgswater_class classifications.

| CASRN | kingdom | superclass | class | subclass |
| --- | --- | --- | --- | --- |
| 7287-19-6 | Organic compounds | Organoheterocyclic compounds | Triazines | 1,3,5-triazines |
| 1031-07-8 | Organic compounds | Organic acids and derivatives | Organic sulfuric acids and derivatives | Sulfuric acid esters |
| 120868-66-8 | Organic compounds | Organoheterocyclic compounds | Pyridines and derivatives | Halopyridines |
| 137-42-8 | Organic compounds | Organic salts | Organic metal salts | Organic alkali metal salts |
| 82-86-0 | Organic compounds | Benzenoids | Naphthalenes | NA |

We use the classified datasets `biosolids_class` and `usgswater_class` in several examples to follow.

If we examine the `usgswater_class` data as follows, we get the contents of

~~~
*R> usgs_class[1:5, c(‘CASRN’, ‘kingdom’, ‘superclass’*,
*+                              ‘class’, ‘subclass’)]*
~~~

Observe in the fifth row, the subclass label is given by NA. This means that the most specific classification label for this chemical is `Naphthalenes` at the `class` level. The chemicals in the previous rows have labels at least down to the `subclass` level, and potentially further, though this table is too small to illustrate the maximum classification depth for all of them (the chemical in the fourth row terminates at the `subclass` level while the three chemicals above it terminate at `level5`, the next level below).

To gain an understanding of these data sets and the chemical classifications they represent, we first look at the number of labels each data set represents, broken down through taxonomic level. The function `label_bars()` takes in a single **data.table** or a (possibly named) list of **data.tables** and constructs histograms of label numbers faceted by data set and by taxonomic level.

~~~
*R> data_list <- list(biosolids_class*,
*+                    usgs_class)*
*R> names(data_list) <- c(‘Biosolids’, ‘USGS Water’)*
*R> label_bars(data_list)*
~~~

From Figure 1 and Figure 2, we see that the distribution of numbers of labels across taxonomic levels is relatively similar for each data set, though in general the `usgswater_class` data has more labels than the `biosolids_class` data at each level. However, this does not give us a sense of how these labels spread through chemical space as defined by ChemOnt nor does it characterize the overlap between the classifications of these data sets. To investigate these aspects of chemical space coverage further, we use tree-based visualizations.

**Figure 1:**
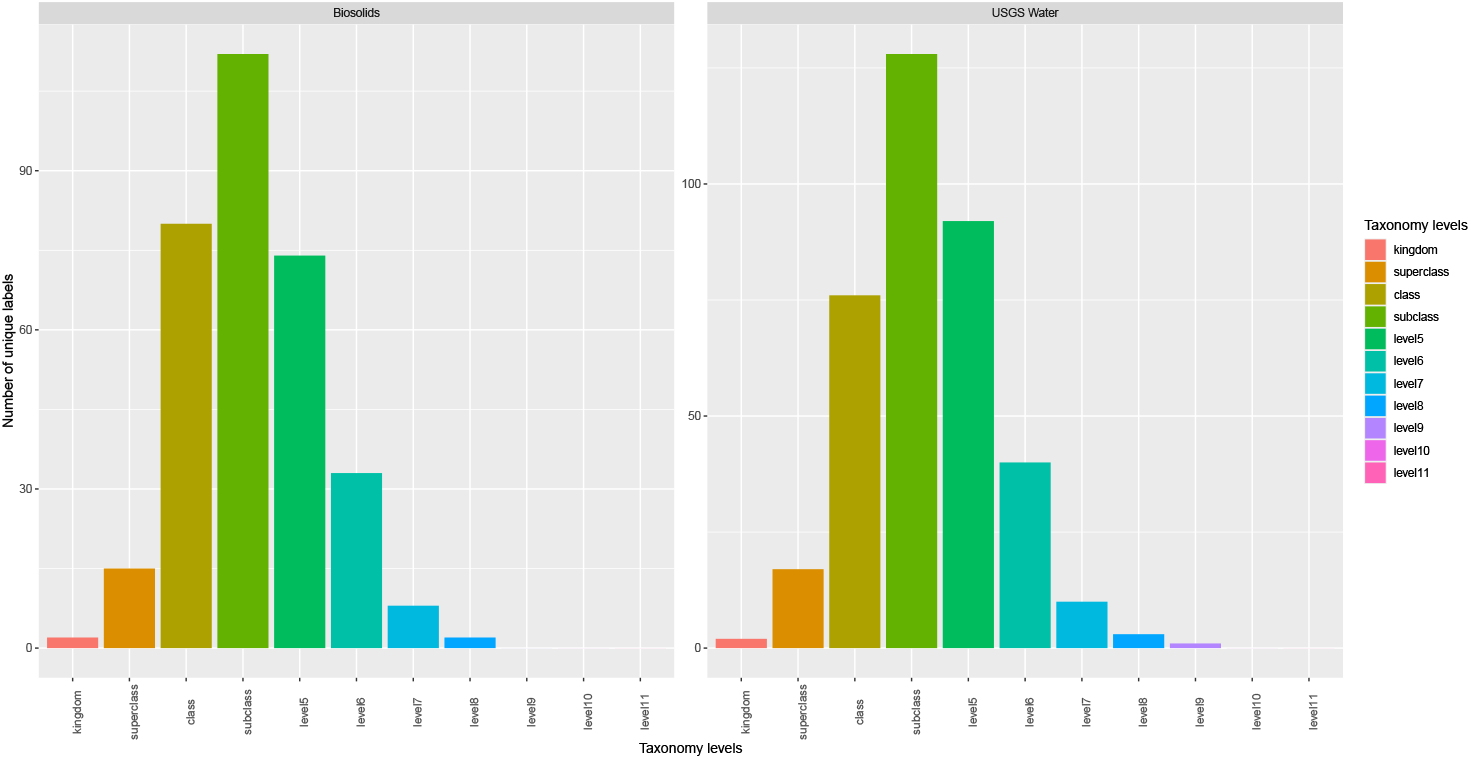
This figure displays the counts of unique labels broken down by each classification level and faceted by dataset.

**Figure 2:**
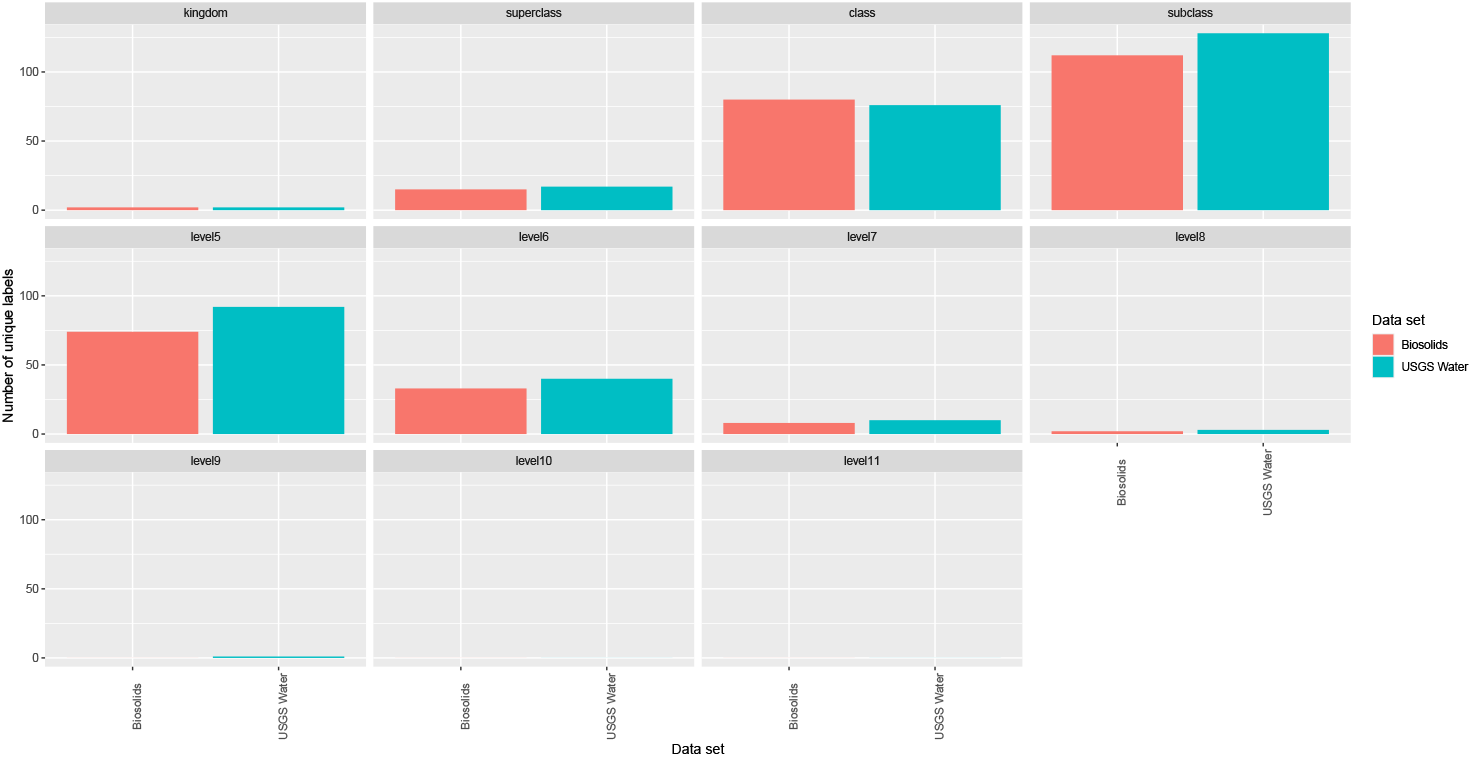
This figure displays the counts of unique labels broken down by dataset and faceted by each classification level.

## 3. Tree visualizations

The tree visualizations developed in **treecompareR** rely heavily on the **ggtree** package, which was designed with visualization of phylogenetic trees in mind. The **ggtree** package extends the widely used **ggplot2** package, greatly broadening the variety and scope of tree visualizations available for use.

We first display the ChemOnt tree, the taxonomy structure of the ChemOnt ontology, in a circular layout format.

~~~
*R> ggtree(chemont_tree) + layout_circular()*
~~~

Figure 3 illustrates the scope of the taxonomy structure of the ChemOnt ontology. From this diagram we create visualizations that highlight the areas of chemical space each data set represents. This ChemOnt tree diagram displays a tree with 3612 tips and 1213 internal nodes (including the root of the tree). As one can easily observe, this creates a dense tree visualization that requires additional attention to be of greater use.

**Figure 3:**
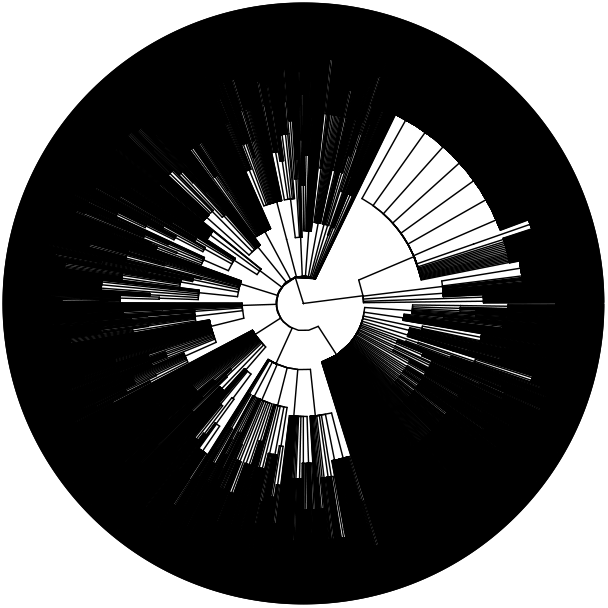
The Chemont tree displayed in a circular layout.

We first examine which parts of this tree are given by classifications of chemicals in each data set. To do this, we use the function `display_subtree()`. This function takes in a **data.table** with classification data and returns a tree diagram highlighting membership for each node and tip of the full ChemOnt tree. Branches and nodes in the tree are highlighted different colors depending on whether they are included in the set of classification labels of the **data.table**. In this example we also display the tip labels.

~~~
*R> display_subtree(data_1 = biosolids_class*,
*+                  name_1 = ‘Biosolids’)*
*R> display_subtree(data_1 = biosolids_class*,
*+                  name_1 = ‘Biosolids’)*
~~~

In the Figure 4 and Figure 5, we observe that the labels and branches represented by classifications from each data set are shaded green while labels and branches not present in the classifications are shaded grey. This helps illustrate where in chemical space these classification labels are located. While it is possible to compare each diagram individually to get a sense of how each data set represents chemical space, it would be better if we could directly compare the subtrees they represent. We can do just that using the exact same function, inputting both data sets into the function `display_subtree()` as the following example demonstrates.

**Figure 4:**
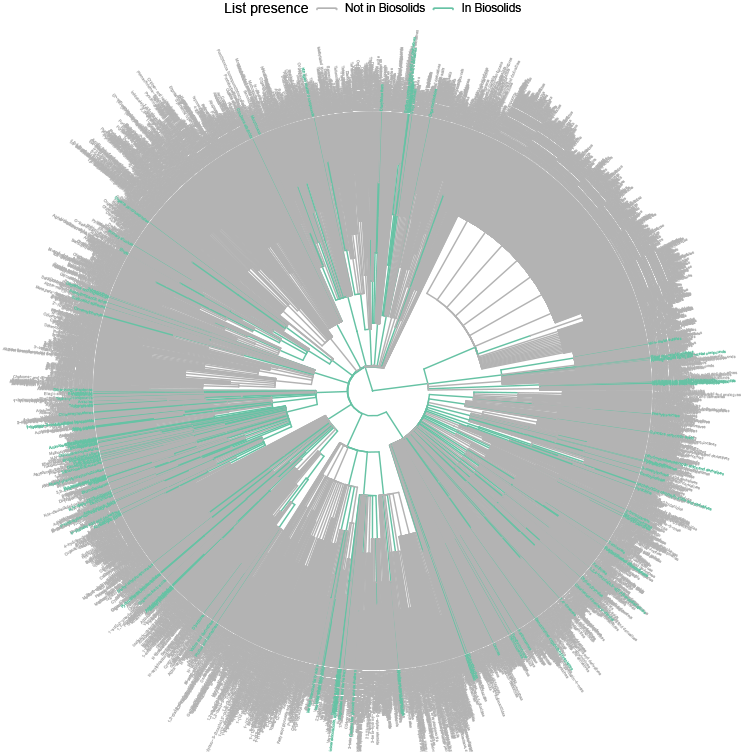
The Biosolids subtree displayed within the full Chemont tree, with tip labels included.

**Figure 5:**
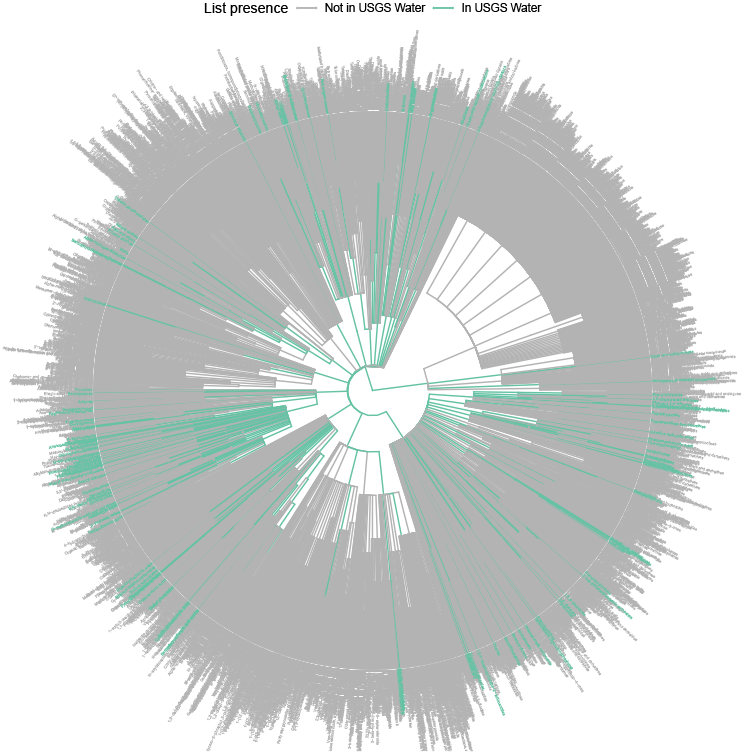
The USGS Water subtree displayed within the full Chemont tree, with tip labels included.

~~~
*R> display_subtree(data_1 = biosolids_class*,
*+                  data_2 = usgs_class*,
*+                  name_1 = ‘Biosolids’*,
*+                  name_2 = ‘USGS Water’)*
~~~

In Figure 6, branches and labels are colored based on whether they are members of classifications of one, both, or neither data set. This gives an indication of how the chemical space represented by each data set overlaps through visualization of the induced subtrees within the full ChemOnt tree. From this visualization, it is also possible to determine the areas in chemical space unique to each data set.

**Figure 6:**
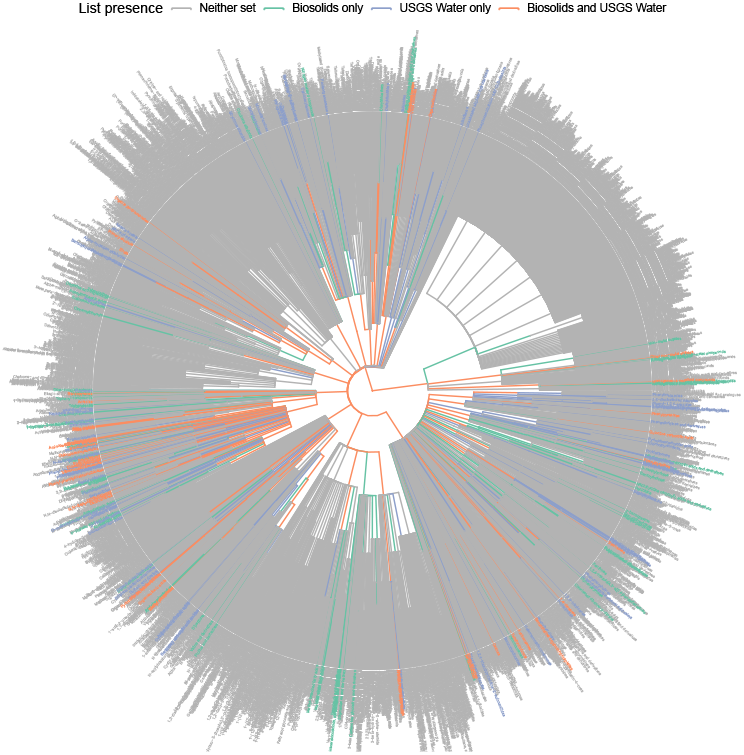
The Biosolids and USGS Water subtrees displayed together within the full Chemont tree, with tip labels included. Classification labels are colored based on whether they are represented by either data set, both, or neither.

While this particular visual can be informative, if the taxonomy is fairly large compared to the subtree induced by a data set, the diagram generated by `display_subtree()` can be cluttered and difficult to interpret. One solution for improving clarity is by pruning away extraneous branches and labels not represented by the subtree. We demonstrate this using the function `prune_and_display_subtree()` on both data sets.

~~~
*R> prune_and_display_subtree(prune_to = biosolids_class)*
*R> prune_and_display_subtree(prune_to = usgs_class)*
~~~

In Figure 7 and Figure 8, the extraneous branches and tips not included in the set of classification labels of the data set are pruned and the remaining subtree is displayed. While there is still a lot of information displayed through the tip labels, this visualization provides a much closer look at the subtree induced by the data set than the previous visual that displayed the subtree within the whole taxonomy. Also observe that the shape of the subtree, while similar to the shape of the subtree displayed within the full ChemOnt tree, may change based on the set of extraneous branches and nodes that are pruned away.

**Figure 7:**
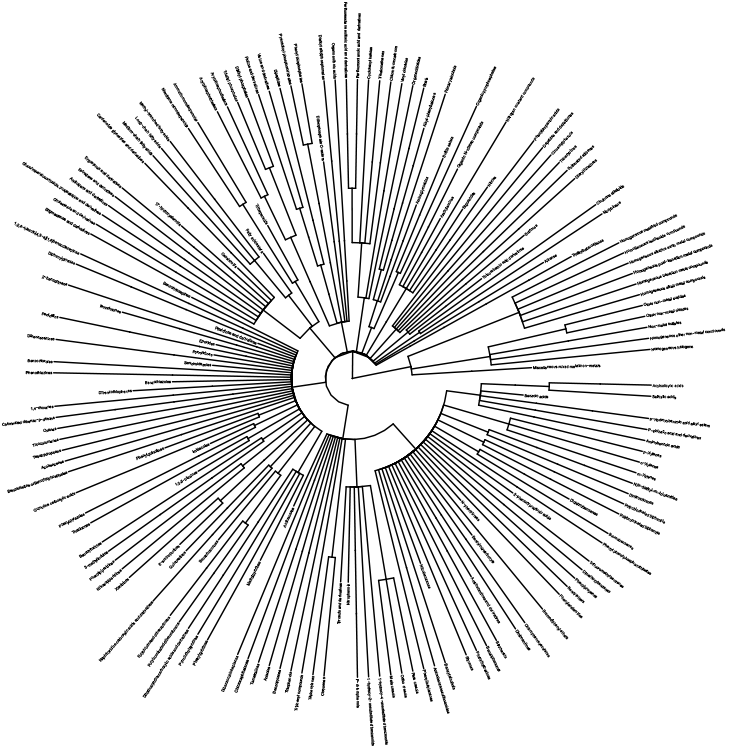
The biosolids subtree pruned from the Chemont tree.

**Figure 8:**
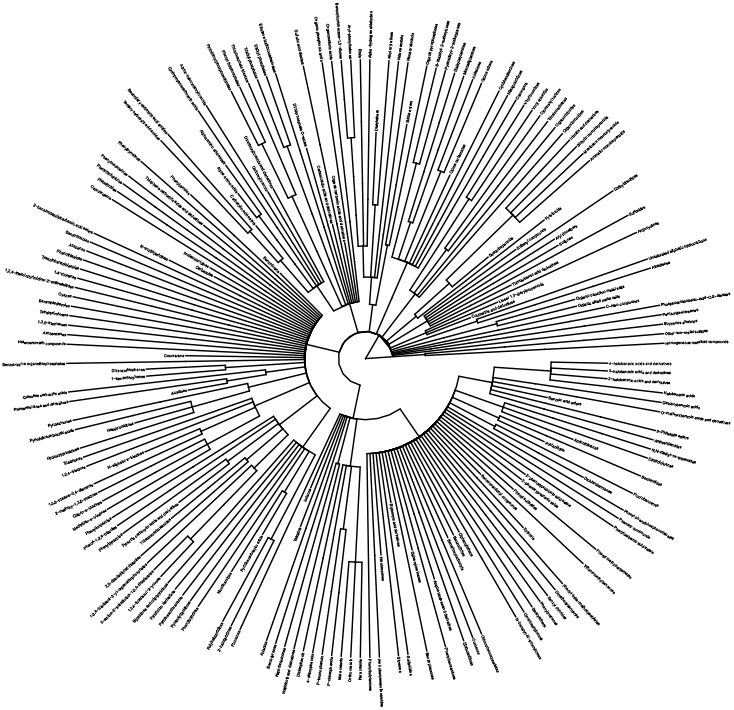
The usgswater subtree pruned from the Chemont tree.

We note a few things about this diagram and the `prune_and_display_subtree()` function. First, the output of the function is a diagram of the subtree (or just the `phylo` object if the function parameter `no_plot = TRUE` is used). Second, while many chemical classifications terminate with a tip label as the most specific label in the classification, there are some classifications that terminate with an internal node. If we were just to delete the internal nodes and tips not represented by classifications for a data set using `ape::drop.tip()`, the internal nodes that were terminal labels (and now are tips of the subtree) for classifications would appear in the diagram somewhere between the outer layer of tips and the root. The `prune_and_display_subtree()` function adjusts the branch lengths to accommodate the new structure of the subtree and extends these newly created tip labels of the subtree to match the pre-existing tip labels. This is recorded in the `edge.length` list of the output `phylo` object. A visual comparison between the branches in the subtree and the subtree within the entire taxonomy may show slight differences as a result of this branch adjustment. Third, with the omission of the extraneous branches and labels, there is the possibility that branches of the pruned subtree have rotate relative to their orientation within the ChemOnt tree. The structure of the subtree is the same, but the visual representation may differ.

In a manner similar to the output of `display_subtree()` when comparing two data sets, we may wish to look at the induced subtrees of two different data sets and compare membership of labels in one subtree with the other subtree. To achieve this, we use the function `data_set_subtrees()` as we illustrate in the following example.

~~~
*R> data_set_subtrees(data_1 = biosolids_class*,
*+                    data_2 = usgs_class*,
*+                    name_1 = ‘Biosolids’*,
*+                    name_2 = ‘USGS water’)*
~~~

Figure 9 and Figure 10 generated by the function `data_set_subtrees()` are useful for illustrating the portions of each data set-induced subtree that are also represented by labels from the classifications of the other data set. The output of the function consists of two tree diagrams, corresponding to the subtrees induced by each data set. Within each diagram, branches and nodes are highlighted different colors depending on whether the associated label is shared by both sets of classifications of the two data sets or is specific to the subtree in which its located. However, much like with the `display_subtree()` output tree visualization, the diagrams generated by `data_set_subtrees()` do not reflect the level to which the overlap occurs, nor do they relate a label in the overlap to the number of chemicals from each data set associated to that classification label.

**Figure 9:**
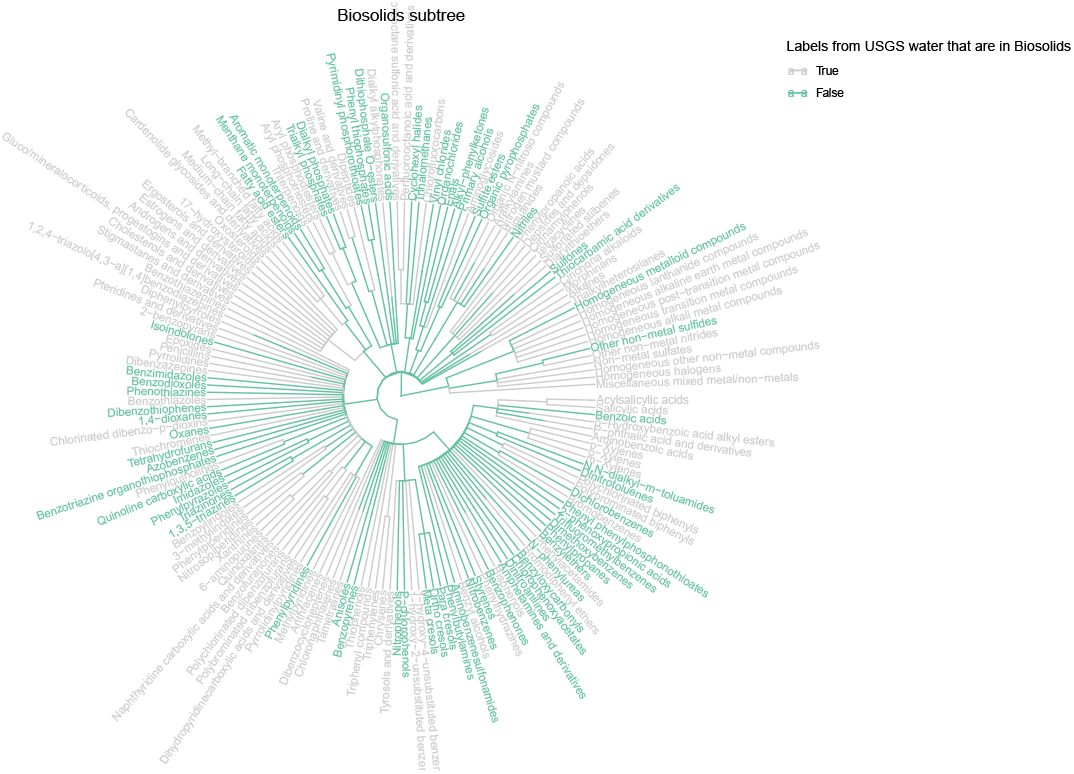
The biosolids subtree pruned from the Chemont tree, with labels colored based on their inclusion in the usgs classification label set.

**Figure 10:**
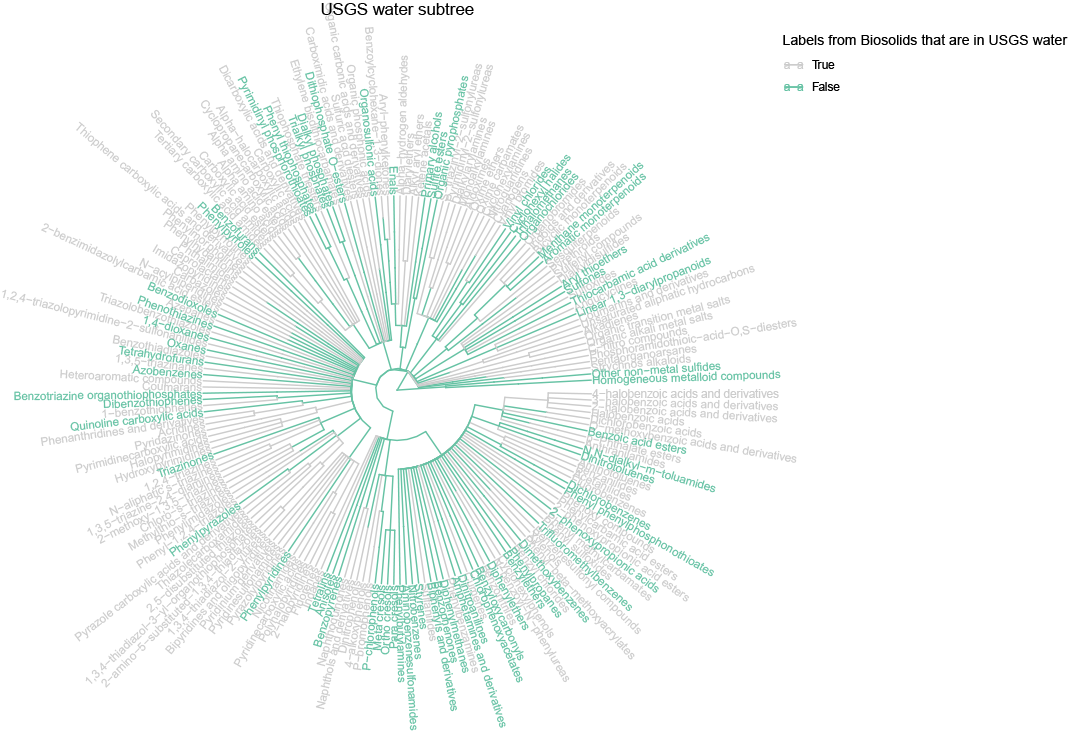
The usgs water subtree pruned from the Chemont tree, with labels colored based on their inclusion in the biosolids classification label set.

To quantify the level of overlap, we need to use a separate tree visualization diagram. We employ the function `leaf_fraction_subtree()` to create diagrams that convey this particular information.

~~~
*R> bio_leaf_frac <- leaf_fraction_subtree(data_1 = biosolids_class*,
*+                                         data_2 = usgs_class*,
*+                                         name_1 = ‘Biosolids’*,
*+                                         name_2 = ‘USGS water’)*
~~~

The output of `leaf_fraction_subtree()` consists of a `data.frame` and a tree visualization. The `data.frame` lists each classification label that shows up as a terminal label in a classification for a chemical in the first data set. In addition, the data frame also displays the number of chemicals that terminate with that label from the first data set, the number that are present in the second data set, and finally the percentage of shared chemicals within the first data set.

The tree visualization features the subtree induced by the first data set. The visualization displays the overlap data contained in the data frame by coloring the terminal label tips based on the degree of overlap. The legend shows the percentage of shared chemicals per terminal classification label using a color gradient that corresponds to the tip and layered colors.

Up to now, the tree visualizations have focused primarily on demonstrating representation within chemical space of classifications of data sets and comparisons between data sets. However, if a data set contains more than just chemical classification data, there may be utility in exploring this additional data. In fact, we can plot additional numeric data from a given data set in layers around the subtree induced by the data set. For instance, in the `biosolids_class` data set, there is a column `AVERAGEṀASS` which gives the average molecular mass for each chemical in the data set. One might be interested in grouping these by their most specific classification label and examining the boxplots of each set of grouped data. We can achieve this by using the function `circ_tree_boxplot()`. Moreover, we can also include taxonomic levels for which labels will be generated in concentric layers.

~~~
*R> circ_tree_boxplot(data = biosolids_class*,
*+                    col = ‘AVERAGE*.*MASS’*,
*+                    title = ‘Biosolids’*,
*+                    tippoint_boxplot = TRUE*,
*+                    layers = c(‘kingdom’, ‘superclass’))*
~~~

One thing to note is that the diagram can be cluttered if we include too many layers. Depending on the size of the data set and scope of its coverage within chemical space, including a few layers corresponding to the middle levels of the taxonomy may result in too many taxonomic labels present in the diagram for reasonable use and interpretation. It is thus important to get a sense of how many labels for each taxonomic level there are in a given data set that one is exploring. For instance, looking at the diagram generated by the function `label_bars()`, one can see that in each data set we have considered, there are over 75 labels in the class taxonomy level alone. Including the class level in the layers parameter of the `circ_tree_boxplot()` function would greatly reduce the clarity of the tree visualizations generated.

## 4. Similarity between trees

The tree visualization functions we have demonstrated thus far have focused on a visual exploration of chemical space and how each data set spreads through it. We have seen how we can examine the overlap in chemical space between data sets and how to examine more closely the subtrees representing each data set. However, there are some more quantitative methods of exploring these overlaps that we have yet to cover.

### 4.1. Similarity measures

The notion of similarity within ontologies and taxonomies has been an area of interest across a variety of disciplines. These aim to compare pairs of nodes in a tree with each other to gain insight on how these nodes relate to each other. Some methods involve examining just the pair of paths from the root of the tree to each of the nodes, while others may examine information more intrinsic to each node. While there is a lot to discuss and explore in the area of similarity measures, we will postpone that to the future and focus on how to use these similarity measures. We will highlight the main points of similarity measures relevant to the uses in this paper and then demonstrate visualizations that leverage them.

Jaccard similarity examines the path from the root to each node in a given pair of nodes, and relates these individual paths to the shared portion of the path (which may possibly be just the root) Pekar and Staab (2002). We can relate these in the following expression

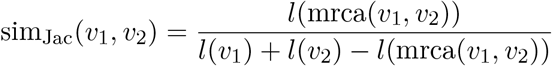

where *l*(*v*_*i*_) denotes the length of the root-to-node path for node *v*_*i*_ and mrca(*v*_*i*_, *v*_*j*_) denotes most recent common ancestor for nodes *v*_*i*_ and *v*_*j*_, the shared node in the root-to-node paths for nodes *v*_*i*_ and *v*_*j*_ furthest from the root (and in some cases possibly the root). Note, the above definition differes slightly in presentation from that in Pekar and Staab (2002), however they are equivalent. This notion of similarity is calculated using the function `general_Jaccard_similarity()`, which takes as parameters a tree and two labels and returns the Jaccard similarity of the two labels within the tree. We exclude the case when the root is one of the nodes being compared. Note, if *v*_*i*_ = *v*_*j*_, then sim_Jac_(*v*_*i*_, *v*_*j*_) = 1.

Some trees may come with a probability distribution, associating to each node a probability such that the sum over all nodes of the probabilities is 1. In the case that a tree does not have such a distribution, we can still derive a node-specific value for each node in the tree. In one such method, we look at the number of descendants a node has and compare this with the total number of nodes in the tree. We would also like for each node to have a higher value than any of its ancestors, with the root having the minimum value over all nodes. One way to achieve this is by comparing the number of nodes in a given subtree with a specified node as the root of the subtree with the total number of nodes of the entire tree, evaluating a monotonically decreasing function on these values at each node, and examining their ratio. In particular, for node *v*_*i*_, define

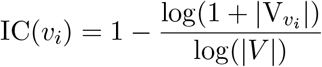

where *V*_*v*_ denotes the descendants of node *v*_*i*_ and *V* is the total number of nodes in the tree, following the treatment by Seco, Veale, and Hayes (2004). In the case that *v*_*i*_ is a tip, it has no descendants so IC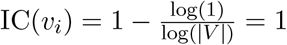. If *v*_*i*_ is the root, then

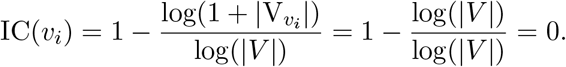

Also note that if *v*_*i*_ is an ancestor of *v*_*j*_, then |*V*_*v_i_*_ > *V*_*v_j_*_| and since the function 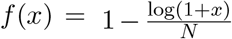 (for *N >* 0) is monotonically decreasing for *x >* 0, it follows that IC(*v*_*i*_) *<* IC(*v*_*j*_). Hence, the function above fits the desired properties.

One way to frame the concept of information content is the idea that the less likely a node appears in a generic path starting from the root and ending at an arbitrary node, the more information it tells when we know the node is on a path given to us. So for instance, the only path a tip can exist on is the unique path connecting it with the root. However, for an ancestor of a tip, there are more paths that include this node, so knowing the ancestor is on a path gives less information about what the whole path might be. Since every path starting at the root includes the root, knowing the root is on the path yields no information.

To generate the information content for a tree, we can use the `attach_information_content()` function which takes in a tree as the only required parameter. This function then adds a data frame to the tree, with each row corresponding to a node of the tree and a column with the information content of each node. The data frame also includes the number of descendants, children, and the level for each node. Note that all of the tips in a tree have 0 descendants and an information content of 1, given by the `log_descendants` column. The root has an information content of 0. Any other nodes have the desired inequality in the values of their information content as it relates to the number of descendants.

Once an information content value is established for each node, we can then use similarity measures that rely on information content to compare nodes.

The first such method is called Resnik similarity, introduced in Resnik (1995). We use the definition given in Seco *et al*. (2004). Resnik similarity is defined for two nodes *v*_*i*_ and *v*_*j*_ as

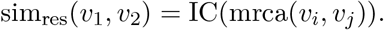

Observe that this is symmetric and when *v*_*i*_ = *v*_*j*_ the similarity measure just returns the information content of the node itself. Note that for nodes that are not tips, their self-similarity is not equal to 1 but instead its associated information content. This implicitly encodes the fact that such a node tells less information than a more specific node does (nodes with fewer descendants). Also, since the information content of the root is zero, any pair of nodes containing the root will return a similarity value of zero.

There are additional similarity measures that build off of Resnik similarity. These will in fact return a value of 1 for self-similarity of nodes unlike in the case of Resnik similarity.

The next method is Lin similarity, introduced in Lin *et al*. (1998) and again use the information content defined in Seco *et al*. (2004). This resembles Jaccard similarity in that it is the ratio of a value derived from the most recent common ancestor of a pair of nodes with values derived from the nodes themselves. In particular, we define

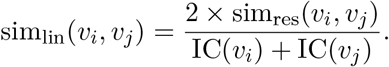

Since the numerator is a scaled form of Resnik similarity, a pair of nodes with the root as their most recent common ancestor will have a value of zero for their Lin similarity. However, unlike Resnik similarity, if *v*_*i*_ = *v*_*j*_, then 2*×* sim_res_(*v*_*i*_, *v*_*i*_) = 2 × IC(*v*_*i*_), so the self-similarity value is equal to 1. Another thing to note is that for a pair of nodes with at least one not equal to the root,

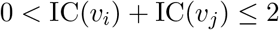

so

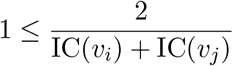

from which it follows that

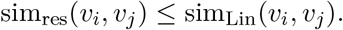

We define the self-similarity of the root to be 1 to reflect the fact that in Lin similarity, self-similarity is equal to 1. This keeps intact the relation above between Resnik similarity and Lin similarity.

An alternative approach to normalizing the similarity value for a pair of nodes takes advantage of the inequality

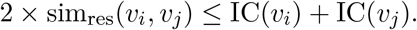

Taking the difference yields the following inequality

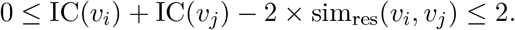

Dividing this inequality by 2 restricts the values the middle expression can take to the interval [0, 1]. However, we want the similarity to be equal to 1 if *v*_*i*_ = *v*_*j*_ and the expression above returns a value of 0 instead. The solution is to take the difference of 1 and this expression, yielding

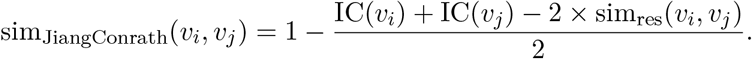

This was developed by Jiang and Conrath in Jiang and Conrath (1997) and is thus named. As with Lin and with Resnik similarities, we use the treatment by Seco *et al*. (2004) for consistency.

Things to note are that when the most recent common ancestor for a pair of nodes *v*_*i*_ and *v*_*j*_ is the root, this similarity measure does not necessarily return a value of zero. In fact, rewriting the expression for Jiang and Conrath similarity, we see that it is equal to

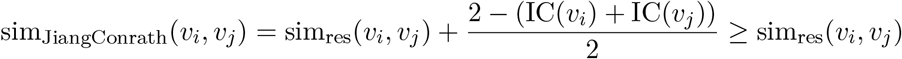

since

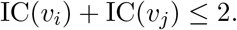

One thing this reflects is that if a pair of such nodes represents a large portion of the tree based on their descendants, they are more likely to be closer to the root and thus more related to each other.

We also observe that

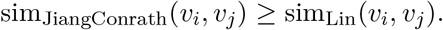

This follows from the following argument. If *v*_*i*_ = *v*_*j*_ are the root,

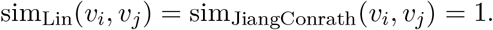

Otherwise, if

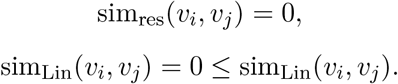

If

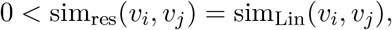

then

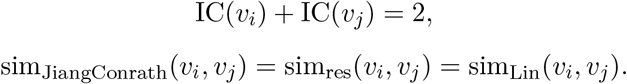

If

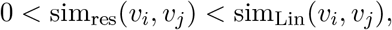

since

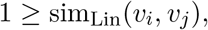

it follows that

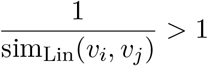

and

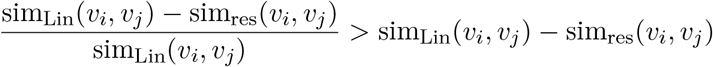

from which it follows that

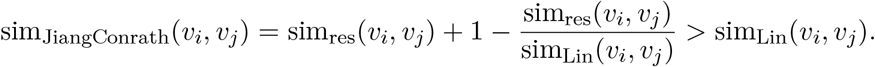

Hence, in all cases

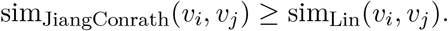

When comparing a pair of nodes that are tips, say *v*_*i*_ and *v*_*j*_, observe that since

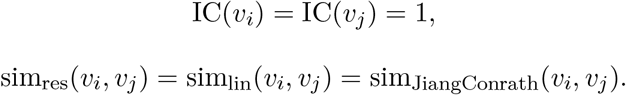

Thus, the three similarity measures all agree on pairs of tips but generally diverge in value otherwise.

Now, the function `attach_information_content()` generates information content for an input tree, and the functions `jaccard_similarity()`, `resnik_similarity()`, `lin_similarity()`, `jiang_conrath_similarity()` all can be used to compare a pair of nodes on a given tree. Moreover, if one wants to compute the similarity for all pairs of nodes in a tree, the function `similarity_matrix()` will do so for a given tree and similarity measure function. The computation is lengthy for the ChemOnt tree, and so these matrices have been pre-computed. They are accessible as `chemont_jaccard`, `chemont_resnik_IC_SVH`, `chemont_lin_IC_SVH`, and `chemont_jiangconrath_IC_SVH`.

### 4.2. Visualizing similarity

Now, we move to using these similarity measures in visualizations. We apply tree visualizations to understand better how data sets relate from the quantitatively-driven point of view of similarity measures. Since similarity measures provide a quantitative approach to exploring the structure of the taxonomy, we can use these numerical relationships to our advantage in studying how data sets cluster relative to one another within chemical space. In particular, we can look at heatmaps generated from a similarity matrix associated to a similarity measure, and where rows and columns are determined by the classification labels representing each data set. From this, we apply hierarchical clustering and examine the portions of each data set corresponding to the individual row and column clusters. This can highlight particularly well the overlapping areas of each data set within chemical space.

First, we construct the heatmap, which requires us to specify a given similarity measure matrix and supply two data sets with classification data. Then we explore clusters more closely using tree diagrams to visualize where the clusters correspond to in chemical space. We can remove extraneous information in this process to get a more precise look at the coverage of both sets within each cluster.

The function `generate_heatmap()` will take in a pair of data sets and compare the induced subtrees using similarity measures. In the example below, we use the Jaccard similarity values of the ChemOnt tree, with rows representing classifications from `usgswater_class` and columns representing classifications from `biosolids_class`. In this diagram, we have specified for there to be nine row and nine column clusters within the hierarchical clustering of the heatmap.

~~~
*R> biosolids_usgs_ht <- generate_heatmap(tree_object = chemont_tree*,
*+                                        matrix = chemont_jaccard*,
*+                                        row_data = USGSWATER_class*,
*+                                        column_data = BIOSOLIDS2021_class*,
*+                                        row_split = 9L, column_split = 9L*,
*+                                        row_title = ‘USGS Water’*,
*+                                        column_title = ‘Biosolids’*,
*+                                        name = ‘Jaccard Similarity’)*
*R> biosolids_usgs_ht*
~~~

In the diagram created by the `generate_heatmap()` function, there are several features that are included. The dendrograms out of which the row and column clusters are determined are displayed on the left side and the top, respectively. Between these dendrograms and the clusters is a layer showing the (logarithm of the) number of times a row or column label occurs within the set of classifications for each data set. To the right of the diagram is a color gradient indicating the similarity values displayed within the heatmap. Observe that in row cluster two and column cluster two, there appears to be a lot of high similarity values. Also note that in row cluster nine, column cluster eight, there are both high similarity values and low similarity values. We analyze these clusters by visualizing the labels associated to each data set in these clusters using tree diagrams.

To visualize the clusters through tree diagrams, we use the function `generate_tree_cluster()`, which takes in a heatmap object and a row and column cluster. It also takes in a tree corresponding to the full taxonomy and a second tree corresponding to a base tree that will be pruned. The function then returns a tree visualization (or pair of them) detailing how each data set contributes to the specific cluster under investigation.

~~~
*R> generate_tree_cluster(tree = chemont_tree, tree_object = chemont_tree*,
*+                        htmap = biosolids_usgs_ht, row_cluster = 2*,
*+                        column_cluster = 2, row_name = ‘USGS Water’*,
*+                        column_name = ‘Biosolids’, isolate_subtree = TRUE)*
~~~

In Figure 14 and Figure 15, corresponding to row cluster two and column cluster two, observe that only the superclass Benzenoids is represented. The first diagram illustrates the portion of the full ChemOnt tree corresponding to the labels in the row and column cluster. The clade defined as all descendants of the Benzenoid label and the label itself are highlighted. in light purple, indicating that the superclass label is shared by classifications in both data sets in this particular cluster.

**Figure 11:**
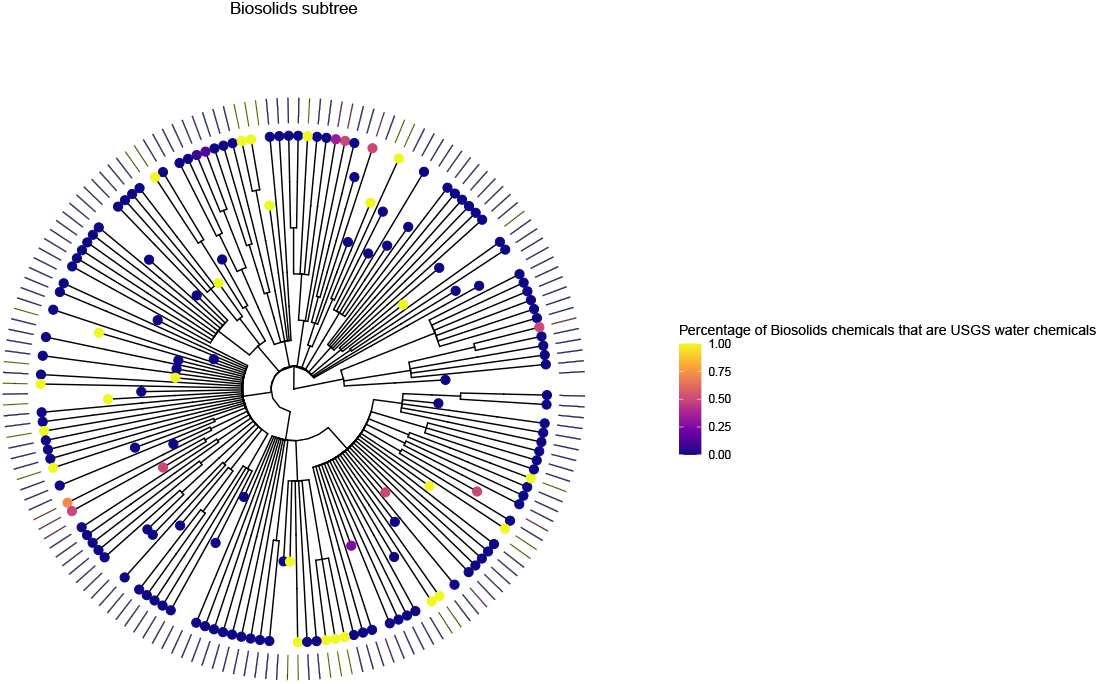
The biosolids subtree with terminal labels colored based on the degree of overlap of chemicals, groups by terminal label.

**Figure 12:**
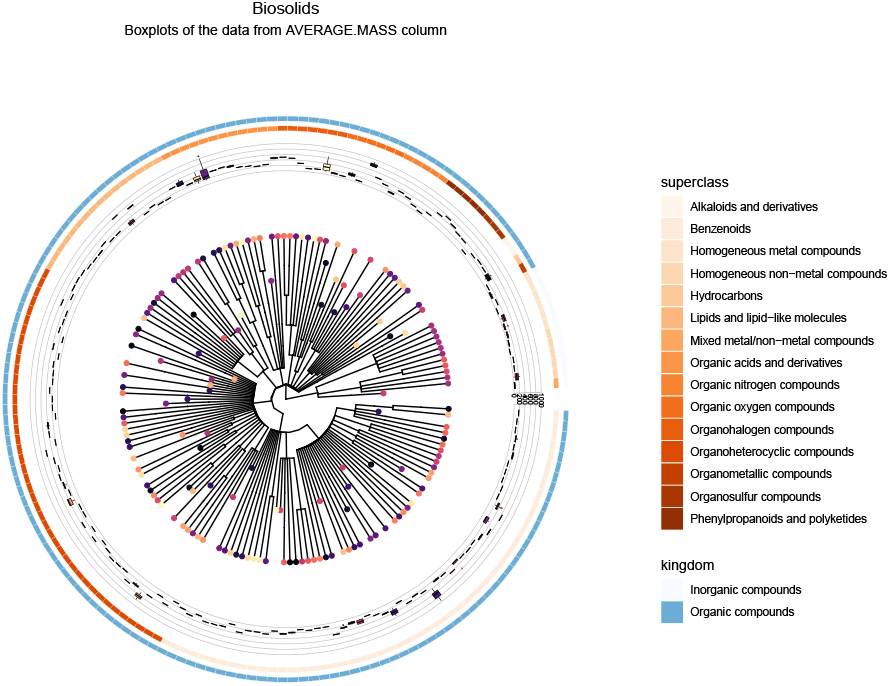
The biosolids subtree with terminal labels colored to match boxplots in outer circular layers. The inner layer displays the distribution of average mass across chemicals, grouped by terminal label. The outer layers specify the `Superclass` and `Kingdom` associated to each terminal label in the classification branches.

**Figure 13:**
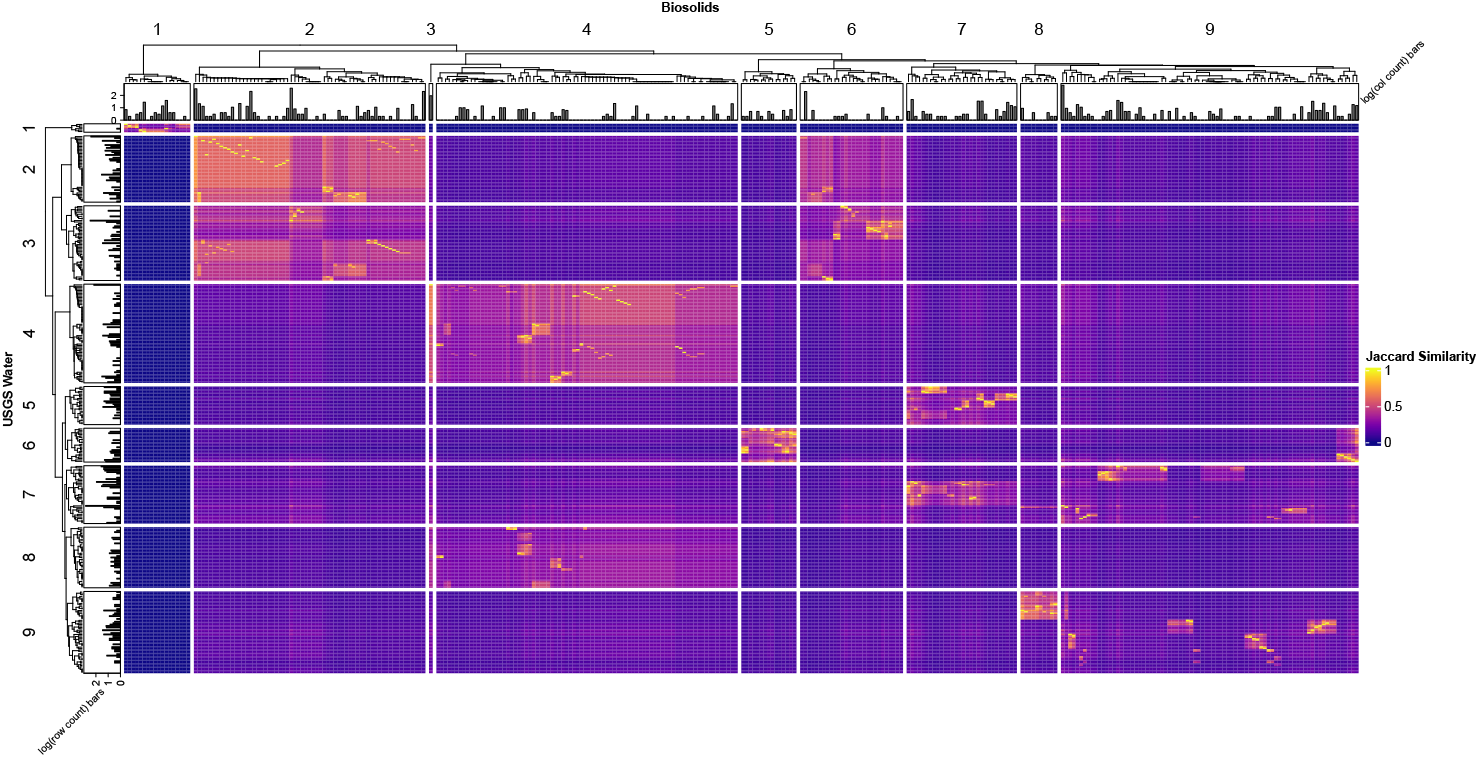
The `ComplexHeatmap` constructed rows corresponding to tip labels from the `USGS water` classification labels and columns corresponding to tip labels from the `Biosolids` classification labels. Clustering is determined based on the corresponding Jaccard similarity measure matrix, appropriately subsetted by rows and columns.

**Figure 14:**
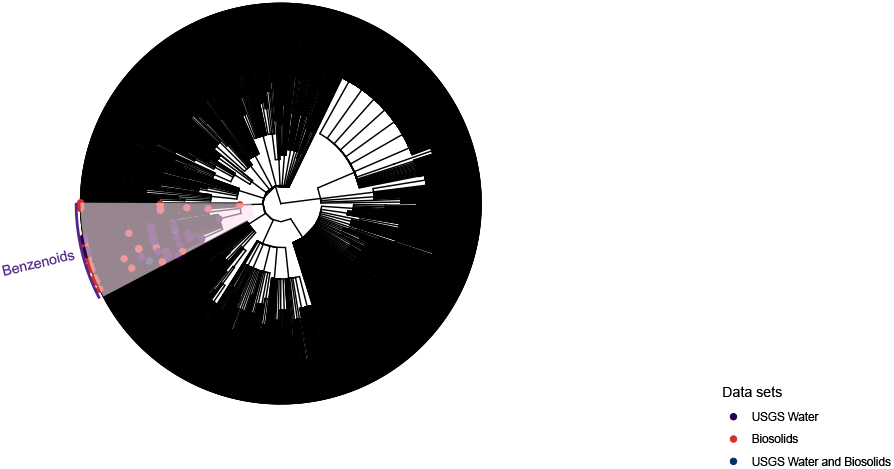
The Superclass represented by row cluster 2 and column cluster 2 within the full ChemOnt tree.

**Figure 15:**
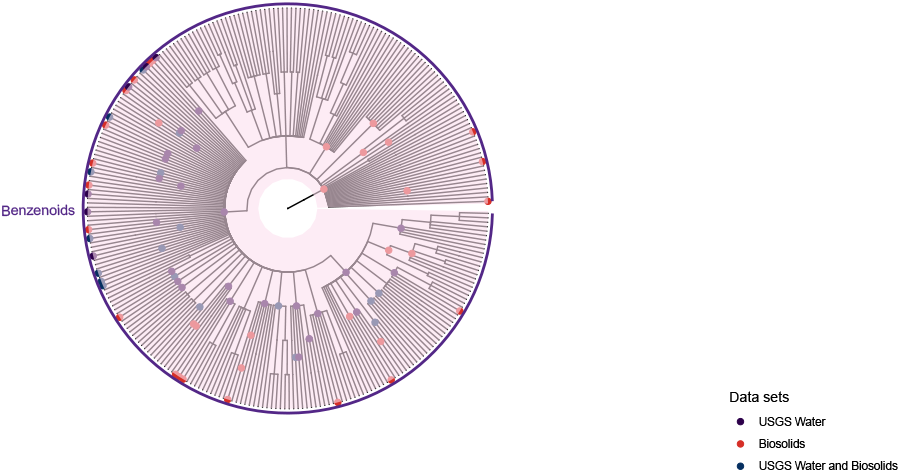
The pruned subtree corresponding to labels in row cluster 2 and column cluster 2. Note the data sets to which each label is associated are noted in the tip label.

In the second diagram, we prune away all extraneous superclass branches, leaving only the Benzenoids superclass. Notice that the labels present in the data sets are indicated by points, colored by whether they are found in one or both data sets. Since the red points greatly outnumber the blue points, one can see that within this cluster the number of labels represented solely by the biosolids data is greater than the number of labels solely represented by the USGS Water data. Also note that there are several purple points, indicating that these labels are shared by both data sets.

~~~
*R> generate_tree_cluster(tree = chemont_tree, tree_object = chemont_tree*,
*+                        htmap = biosolids_usgs_ht, row_cluster = 9*,
*+                        column_cluster = 8, row_name = ‘USGS Water’*,
*+                        column_name = ‘Biosolids’, isolate_subtree = TRUE)*
~~~

In the pair of trees in Figure 16 and Figure 17, corresponding to row cluster nine and column cluster eight, we observe that there are several superclasses represented, seven corresponding to just USGS classifications (highlighted in light green) and one shared by both Biosolids and USGS (highlighted in light purple). Consequently, there are labels from the USGS Water data set present in all of the superclass clades that are highlighted, though the labels from the biosolids data set are only found within the Organic nitrogen compounds superclass.

**Figure 16:**
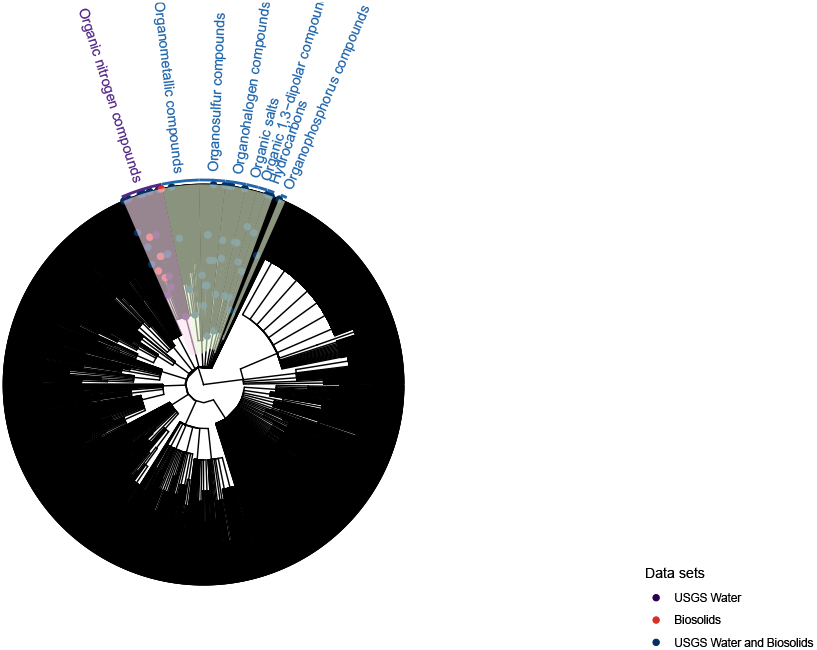
The Superclasses represented by row cluster 9 and column cluster 8 within the full ChemOnt tree.

**Figure 17:**
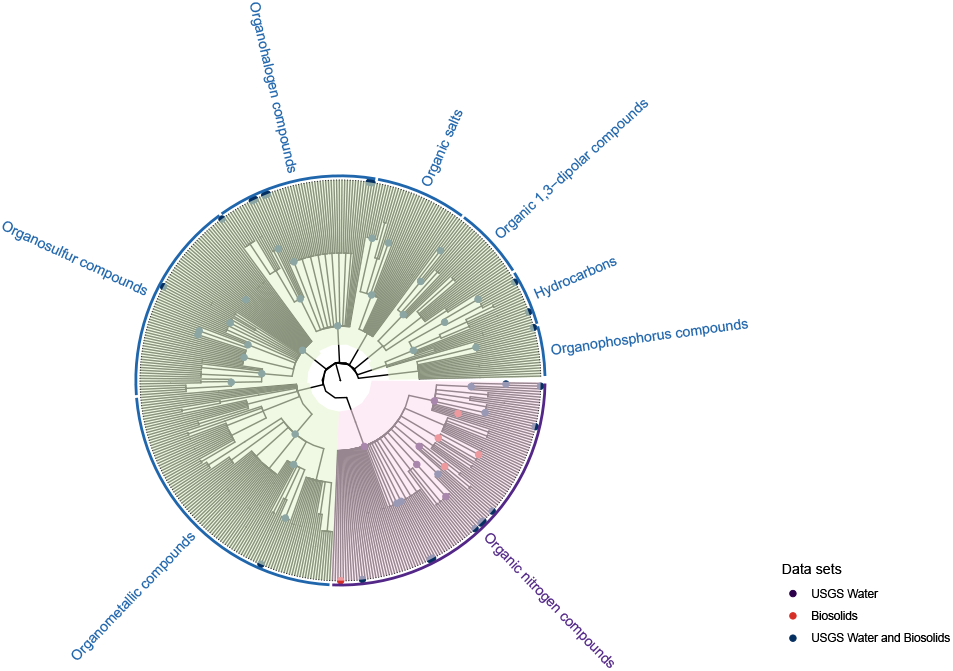
The pruned subtree corresponding to labels in row cluster 9 and column cluster 8. Note the data sets to which each label is associated are noted in the tip label.

In the Figure 17, we can observe this more closely. Within the shared clade given by the Organic nitrogen compounds clade, there are very few labels that are just given by the biosolids classifications, while most labels are solely from the USGS Water classifications and some are shared by both data sets.

## 5. Summary and discussion

The visualization tools provided by **treecompareR** allow for an analysis of chemical space not previously available. This can benefit researchers analyzing data sets in a variety of ways. While there are already ‘chemical fingerprints’ in use for characterizing chemical structure, these fingerprints can be hard to interpret especially within a large data set or for those without specialized knowledge of those chemical structures. The ClassyFire framework implicitly encodes some of this information into the taxonomic structure of the ontology and provides a classification scheme more accessible to a broader group of users. However, visual interpretation of this data is an important aspect of the data analysis in addition to quantitative interpretations, and **treecompareR** provides tools to address both of these needs. This allows users an increased ability to answer questions that naturally arise concerning domain of applicability of models, to identify data gaps that require additional data gathering efforts, First, using the function `label_bars()`, one can make a preliminary analysis on the broad coverage of labels within chemical space that one or several data sets produce. The number of labels per taxonomic level can tell the user whether a data set is confined to a few labels at a specific taxonomic level or spreads through a greater portion of the labels in the given level. For instance, in Figure 2, we observe that USGS Water covers more labels than Biosolids, but that the difference is relatively small.

Next, we turn to the tree diagrams to gain a visual understanding of how the data sets overlap within chemical space. By plotting two subtrees induced from data sets together, we can observe whether the data correspond to the same areas of chemical space or are in distant areas. For example, in Figure 6, we observe that there is indeed a decent amount of overlap. These data, while representing some areas of chemical space specific to one data set or the other, have a non-trivial amount of overlap. This gives some evidence to suggest that at least in the areas of overlap, models trained on one data set would have a reasonable likelihood of being applicable for use with the other data set.

To study the overlap further, one can use either the function `data_set_subtrees()` or the function `leaf_fraction_subtree()` for a more in-depth examination of how the data sets relate. These allow for a simple visual examination of the overlap in each induced subtree as well as a quantitative examination of the degree of the overlap. In Figure 9 and Figure 10, we can get a clear idea of what classifications are shared by each data set and areas specific to one or the other. To quantify this overlap, we perform an examination of the tree diagram in Figure 11 and of the associated data.frame. There are 80 taxonomic labels shared between the two subtrees induced by the biosolids and usgswater data sets. This accounts for 40.4% of the Biosolids subtree (80/198) and 34.63% of the USGS Water subtree (80/231). Moreover, there are 44 classification labels associated with 65 chemicals shared between the data sets. These 44 classification labels are the most specific labels in the classifications of the associated chemicals, the ‘terminal labels’ in the classification data of each classified chemical. For the 65 chemicals that are shared between the data sets, it would be useful to explore further the concentrations of there occurrence in both water and biosolids. If chemicals expected to partition to solids are still found at high concentrations within water, this would suggest that there is a high background concentration of these chemicals. On the other hand, if the concentration within water were relatively low, it might suggest that these chemicals are removed effectively by wastewater treatment processes and do not persist at great levels within the environment after water treatment.

For a more quantitative approach to identifying overlaps of chemical space coverage from each data set, we turn to similarity measures. By examining the heatmap in Figure 13, we can identify clusters of high similarity and clusters of low similarity. For the purposes of training a model and studying the associated domain of applicability, it is reasonable to expect that chemicals with classifications grouped within clusters of high similarity values between the training set and target set may have a greater likelihood of falling within the model domain of applicability. For example, an examination on row cluster two and column cluster two, featured in Figure 14 and Figure 15, leads one to conclude that chemicals with a superclass label of ‘Benzenoids’ may have more reliable model predictions when using the `biosolids_class` data to train a model. Likewise, chemicals with a superclass label of ‘Organic nitrogen compounds’ would also have a more reliable model prediction due to the overlap featured in row cluster nine and column cluster eight, and displayed in Figure 16 and Figure 17. In fact, since most rows in Figure 13 have a corresponding row cluster of higher similarity, it is reasonable to expect that many of chemicals contained in the `usgswater_class` data set would have reliable model predictions produced from a model that was trained on the `biosolids_class` data set; the `usgswater_class` data set is likely well covered by the applicability domain of a model trained using the `biosolids_class` data.

## 6. Conclusion

The `R` package **treecompareR** provides a suite of tools for visualizing hierarchical chemical classification data associated to ClassyFire and for determining the similarity of subtrees of the ChemOnt tree induced by chemical data sets. The visualization functions provide tools for a qualitative analysis of data set similarity and overlap, while the similarity functions provide a more quantitative approach. With both of these sets of tools, researchers can explore questions concerning the overlap of distinct data sets and how that might inform the training and testing of models that use chemical data. Specifically, these can elucidate potential data gaps in the domain of applicability of a model or suggest when a data set might be suitable for use with a trained model. We demonstrate these ideas of visual and quantitative exploration of two real data sets available from the CompTox Chemicals Dashboard. While this showcases many opportunities to explore these data sets, the functionality provided by **treecompareR** is not meant as a replacement of existing techniques, but rather to augment the approaches researchers might take to characterize their data.

## Computational details

The results in this paper were obtained using `R` 4.4.1 with the **treecompareR** 1.0.0 package, available on GitHub at https://github.com/USEPA/treecompareR. `R` itself and all other packages used are available from the Comprehensive `R` Archive Network (CRAN) at https://CRAN.R-project.org/, BioConductor at https://www.bioconductor.org/ or from GitHub.

## Acknowledgments

The authors would like to acknowledge the support of colleagues within the US EPA Center for Computational Toxicology and Exposure. In addition, this project was supported in part by an appointment to the Research Participation Program at the Center for Computational Toxicology and Exposure, Office of Research and Development, U.S. Environmental Protection Agency, administered by the Oak Ridge Institute for Science and Education through an interagency agreement between the U.S. Department of Energy and EPA.

## Disclaimer and Conflict of Interest Statement

The views expressed in this publication are those of the authors and do not necessarily represent the views for policies of the U.S. Environmental Protection Agency. Reference to commercial products or services does not constitute endorsement. The authors declare no conflict of interest.

